# Membrane-bound Interleukin-1α mediates leukocyte adhesion during atherogenesis

**DOI:** 10.1101/2023.05.31.543077

**Authors:** Christina Mäder, Thimoteus Speer, Angela Wirth, Jes-Niels Boeckel, Sameen Fatima, Khurrum Shahzad, Marc Freichel, Ulrich Laufs, Susanne Gaul

## Abstract

**Background:** The interleukin-1 (IL-1) family and the NLR family pyrin domain-containing 3 (NLRP3) inflammasome contribute to atherogenesis but the underlying mechanism are incompletely understood. Unlike IL-1β, IL-1α is not dependent on the NLRP3 inflammasome to exert its pro-inflammatory effects. Here, a non-genetic model was applied to characterize the role of IL-1α, IL-1β and NLRP3 for the pathogenesis of atherosclerosis.

**Methods:** Atherogenesis was induced by gain-of-function PCSK9-AAV8 mutant viruses and feeding of a high-fat western diet (WTD) for 12 weeks in C57Bl6/J wildtype mice (control) and in Il1a^-/-^, Nlrp3^-/-^, and Il1b^-/-^ mice.

**Results:** Il1a^-/-^ mice showed reduced atherosclerotic plaque area in the aortic root with lower lipid accumulation, while no difference was observed between wildtype, Nlrp3^-/-^ and Il1b^-/-^ mice. Serum proteomic analysis showed a reduction of pro-inflammatory cytokines (e.g. IL-1β, IL-6) in Il1a^-/-^ as well as in Nlrp3^-/-^ and Il1b^-/-^ mice. Bone marrow dendritic cells (BMDC) of WT, Nlrp3^-/-^ and Il1b^-/-^ mice and primary human monocytes showed translocation of IL-1α to the plasma membrane (csIL-1α) upon stimulation with LPS. The translocation of IL-1α to the cell surface was regulated by myristoylation and increased in mice with hypercholesterolemia. CsIL-1α and IL1R1 protein-protein interaction on endothelial cells induced VCAM1 expression and monocyte adhesion, which was abrogated by the administration of neutralizing antibodies against IL-1α and IL1R1.

**Conclusions:** Il1a^-/-^ mice, but not Nlrp3^-/-^ or Il1b^-/-^ mice, are protected from atherosclerosis after induction of hypercholesterolemia independent of circulating cytokines. Myristoylation and translocation of IL-1α to the cell surface in myeloid cells facilitates leukocyte adhesion and contributes to the development of atherosclerosis.

Graphical abstract.
The role of cell-surface (cs) IL-1a in the initiation of atherosclerosis.

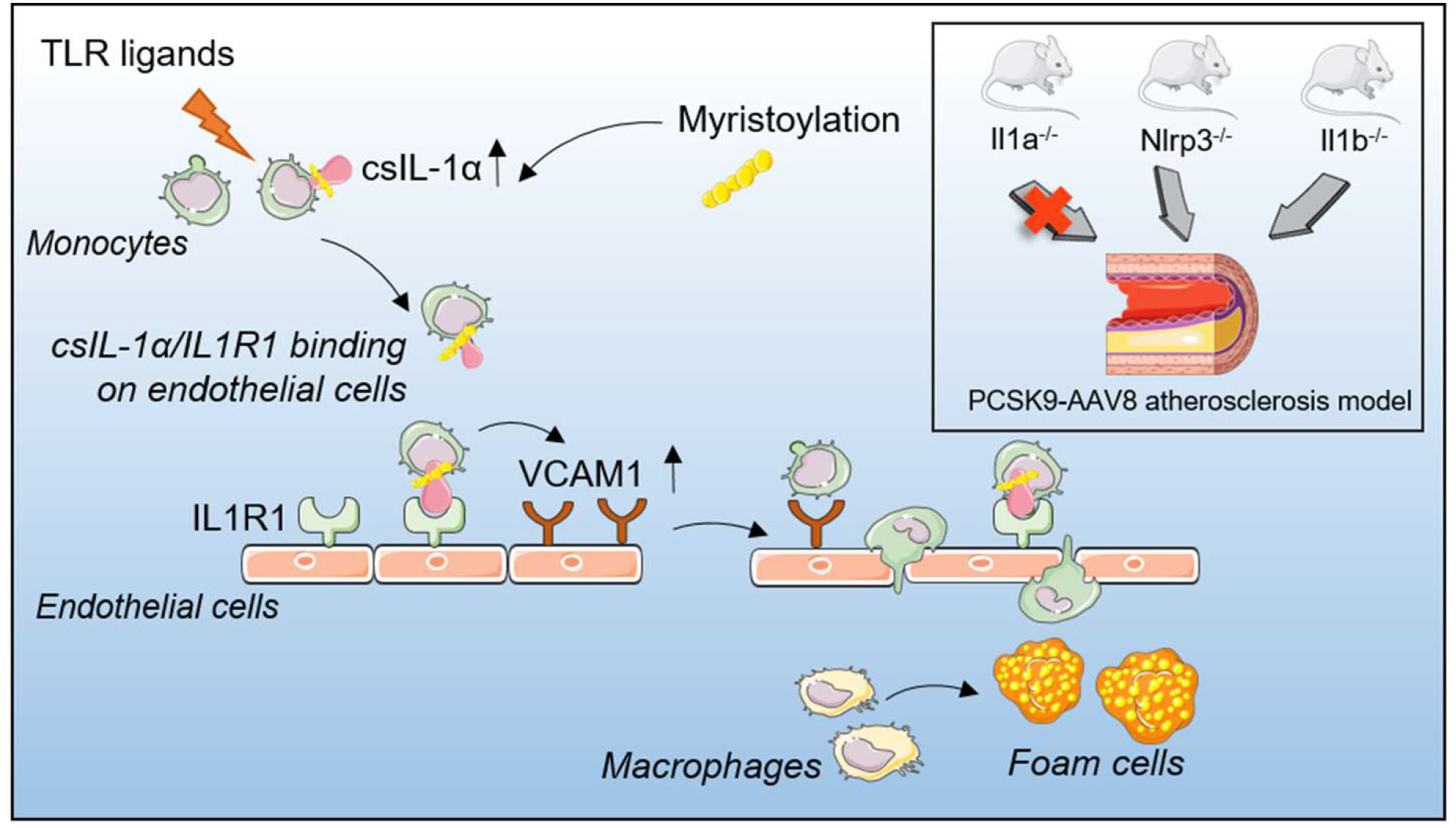

## Introduction

Atherosclerosis is a chronic inflammatory disease driven by intimal lipid accumulation ^1,2^. Reactive oxygen species modify plasma lipoproteins in the subendothelial space, where they are engulfed and digested by macrophages ^3^. Processing of oxidized lipoproteins inside the cell leads to cholesterol crystal formation and activation of the NOD-, LRR- and pyrin domain-containing protein 3 (Nlrp3) inflammasome, initiating the release of pro-inflammatory cytokines such as IL-1β ^4,5^. During plaque progression, different subsets of leukocytes adhere to endothelial cells and infiltrate the atherosclerotic plaque contributing to the inflammatory environment ^6,7^.

IL-1 cytokines are considered inflammatory mediators of atherosclerosis ^8^. Two related but distinct IL-1 genes, *IL1A,* and *IL1B,* encode IL-1α and IL-1β, respectively. Both share the same receptor, and upon binding of either cytokine, interleukin 1 receptor type I (IL1R1) associates with interleukin 1 receptor accessory protein (IL1RAP/IL1R3) to form a trimeric structure that mediates NfκB-dependent pro-inflammatory signaling ^9^. The inactive IL-1β precursor protein is cleaved by caspase 1 to the mature form of IL-1β ^10^.

In contrast to IL-1β, both the precursor and the cleaved form of IL-1α are biologically active. Pro-IL-1α can also be presented on the cell surface (csIL-1α) upon activation of the TLR, e.g., in the presence of lipopolysaccharides (LPS) ^11^. The precise mechanism of binding IL-1α to the outer cell membrane is incompletely understood ^12^. It was observed that csIL-1α plays a role in acute myocardial infarction (AMI) and chronic kidney disease (CKD) and is associated with an increased risk of cardiovascular events ^13^. However, its functional role in atherogenesis is not fully elucidated.

Previous studies of atherogenesis in mice depended on cross-breeding with either Ldlr- or ApoE-deficient animals. Both models exhibit high plasma cholesterol levels and the development of atherosclerotic lesions in susceptible areas ^14^. Studies on genes involved in atherosclerosis require inbreeding in either Ldlr^-/-^ or ApoE^-/-^ animals with the associated potential confounders. However, studies of gene candidates without the genetic bias of the atherosclerosis model itself were not available. We therefore applied a novel nongenetic gain-of-function PCSK9-AAV8 atherosclerosis model to address the question whether atherosclerosis development depends on IL-1α or the Nlrp3 inflammasome and whether IL-1α expressed at the cell surface is involved in the development of atherosclerosis ^15^.

## Material and Methods

### Mice

WT C57Bl6/J (n=10) and Nlrp3^-/-^ (B6.129S6-Nlrp1btm1Bhk/J, n=5) were purchased from Charles River (Sulzfeld, Germany). Il1a^-/-^ (n=6) and Il1b^-/-^ (n=5) mice were a kind gift from Prof. Manfred Kopf (Eidgenössisch Technische Hochschule Zurich, Zürich, Switzerland). To induce atherosclerosis, the mice were injected with a mutated AAV8 virus, as described previously ^15^. Briefly, age-matched littermates were injected with rAAV8-PCSK9^D377Y^ (1.0×10^11^ viral genomes/mouse) or Saline as control at 10 weeks (Figure 1A). Mice received a western high-fat diet (Ssniff, 21 % fat, 0.21 % cholesterol) one-week post-virus injection for 12 weeks. Saline-treated control mice were fed a normal chow diet. Body weights were measured weekly. Mice were sacrificed after the given time point, and organs were harvested. In brief, the mice were anesthetized with 4-5% isoflurane. The absence of the pedal reflex confirmed successful anesthesia. Afterward, blood was collected, and mice were perfused for organ harvest. All protocols were approved by the Institutional Animal Care (42/2019, Veterinary Office of the Saarland) and were consistent with the guidelines from Directive 2010/63/EU of the European Parliament.

**Figure 1:**
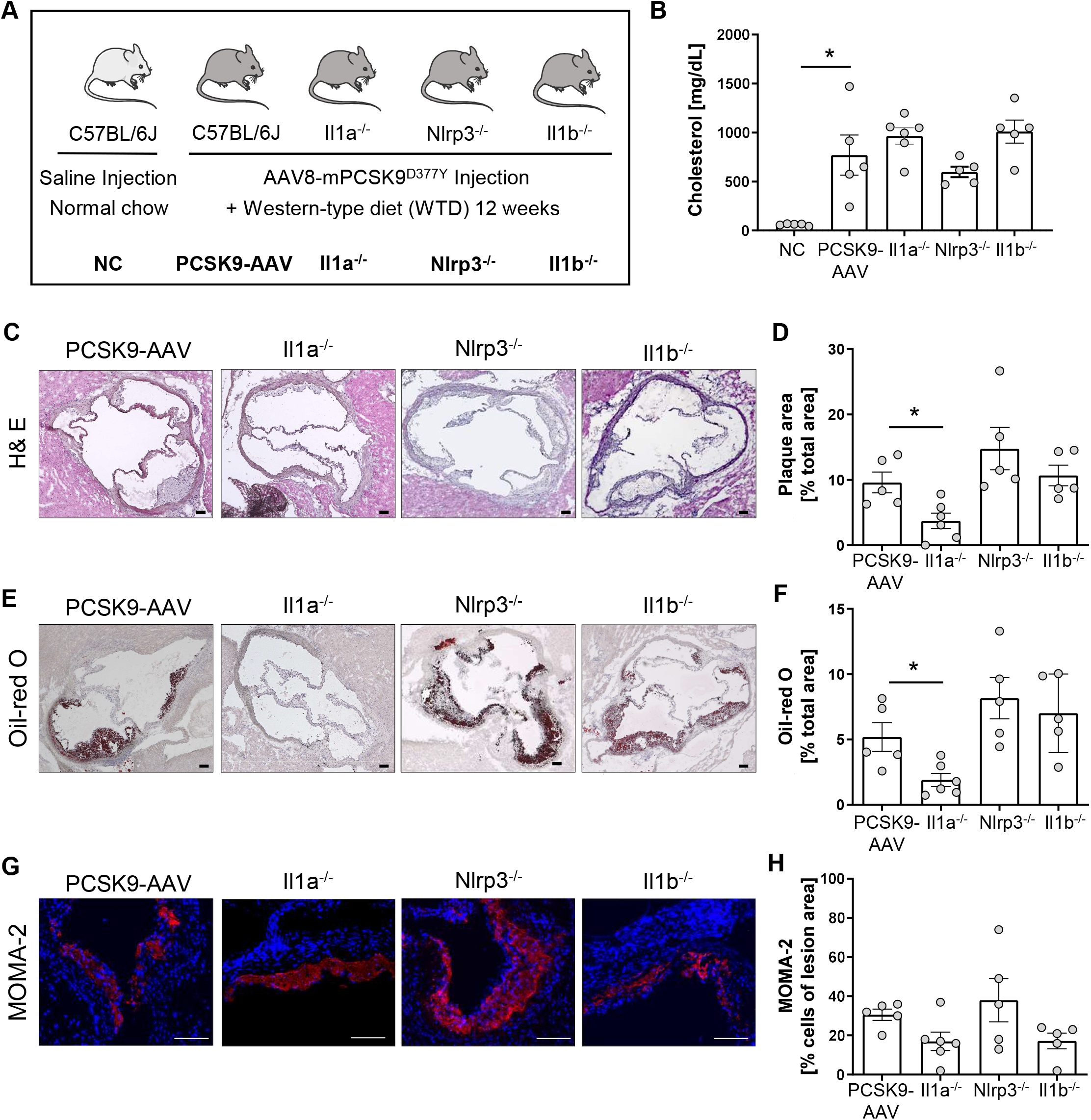
Il1a knockout mice are protected from hyperlipidemic-induced atherosclerosis independent of the NLRP3-inflammasome. **A:** Schematic overview of the experimental groups. Mice were injected with either rAAV8-PCSK9^D377Y^ or Saline. One-week post-injection, mice were fed a western-type diet (21 % fat, 0.21 % cholesterol) for 12 weeks. Saline-injected mice received a normal chow (NC) diet. **B**: Bar graph of plasma cholesterol [mg/dL] after 12 weeks of diet intervention. **C-H:** Representative histological images of the aortic root stained with Hematoxylin/Eosin (H& E, **C**), Oil-red O (ORO, **E**), and monocyte and macrophage (MOMA-2) staining (**G**) with their corresponding bar graphs (**D, F, H**). Data are presented as mean ± SEM. One-way ANOVA was performed with Sidak’s multiple comparisons posthoc test (*p < 0.05). H& E and ORO stainings were imaged with 4 × magnification, scale bar 100 µm (**C, E**), MOMA-2 staining with 10 × magnification, scale bar 100 µm (**G**). Data are presented as mean± SEM of PCSK9-AAV8 (n=5), Il1a^-/-^ (n=6), Nlrp3^-/-^ (n=5), and Il1b^-/-^ (n=5) animals.

### Histology

Characterization of atherosclerotic lesions was performed in the aortic root. Cryopreserved sections (10 µm) were used for Hematoxylin and Eosin (H& E), MOMA-2, and Oil-red O (ORO) staining. MOMA-2 and ORO staining were performed as described previously ^15^. H& E staining was performed by fixation of cryoslides with Xylene 3x 5 min, followed by a descending series with alcohol. Slides were washed with H2O before the hematoxylin staining for 4 min. Then, sections were washed with H2O and incubated with Scott’s bluing water for 20 seconds, followed by 1 min in 95 % EtOH. Staining with Eosin was performed for 1 min, and sections were incubated in an ascending series of alcohol followed by 3x 5 min in xylene. Stained sections were then covered with xylene-based mounting solution and coverslips. ImageJ software (Version 1.53k) was used for image analysis. MOMA-2 staining was imaged with a ZEISS Elyra microscope. The percentage of MOMA-2 positive area per lesion was calculated by determining the MOMA2 positive area and dividing it by the total plaque area.

### Isolation and culture of murine Bone Marrow-Derived Dendritic Cells (BMDC)

Bone marrow cells were isolated from wildtype mice ^16^. 2×10^6^ bone marrow cells were plated on a 10-cm dish in RPMI-1640 medium (Sigma Aldrich, USA) containing 10 % fetal calf serum (Gibco, Thermo Fisher Scientific, USA), 1% Penicillin-Streptomycin, and 50 mM β-Mercaptoethanol (Gibco, Thermo Fisher Scientific, USA). To induce dendritic cell differentiation, the medium was supplemented with 20 ng/ml GM-CSF (Preprotech, Thermo Fisher Scientific, USA) and 10 ng/ml IL-4 (Preprotech, Thermo Fisher Scientific, USA) ^17^. Cells were harvested on day 7 by collecting the suspension cells in the supernatant. Differentiation was confirmed using flow cytometry by detecting the MHC-II and Cd11c double-positive cells (min. 70% double-positive cells) (Figure S2A). Harvested cells were replated for further experiments in a fresh culture dish.

### Isolation and stimulation of human primary monocytes

Human primary monocytes were isolated from buffy coats of healthy volunteers via Ficoll (Cytiva, USA) gradient centrifugation (ethics license 272-12-13082012). Isolated PBMCs were incubated with CD14 beads (Milteny Biotech, Germany) for 15 min at 4°C and isolated with MACS magnetic separator. Isolated monocytes were cultured in RPMI-1640 (Sigma Aldrich, USA) with 10% Fetal calf serum and 10 mM Sodium Pyruvate (Merck, Germany). Cells were stimulated with 100 ng/ml ultrapure LPS (upLPS, Invivogen, USA) overnight with or without pre-incubation of 10 µM/ml NLRP3-Inhibitor MCC-950 (Invivogen, USA) for 1h.

### Culture of Human Umbilical Vein Endothelial Cells (HUVEC)

Human Umbilical Vein Cells (HUVECs) were cultured in EBM Endothelial Cell Growth Basal Medium (Lonza, Swiss) supplemented with EGMTM Endothelial Cell Growth Medium SingleQuots (Lonza, Swiss). Passages 4 to 8 were used for experiments.

### Serum analysis

Serum was collected for the measurement of plasma lipids and plasma proteins. 500-700 µl of blood were collected from each mouse and centrifuged at 3.500× g for 10 min at room temperature. Blood serum was transferred to a fresh 1.5 ml tube and snap-frozen in liquid nitrogen. Cholesterol was measured using LabAssay Cholesterol (Fuji Film, Japan) following the manufacturer’s instructions.

The Olink Mouse Exploratory Panel was used to measure circulating proteins. Serum was prepared according to the manufacturer’s instructions (Olink, Sweden) and measured with the Mouse Exploratory 96 panel. Protein concentration was determined using the proximity extension assay (PEA) technology described elsewhere^18^. Briefly, oligonucleotide-labeled antibodies bind to their target proteins where they come into proximity with other labeled antibodies that bind to the same target. The oligonucleotides hybridize in proximity and form a basis for qPCR and quantification. The number of qPCR cycles stands in relation to the protein concentration and gives the arbitrary unit normalized protein expression (“NPX”) as the readout. To examine changes in the secretome in the hyperlipidemic mouse model vs. mice fed a normal chow (NC) diet, two-sideded t-test was performed, followed by enrichment analysis of the significant regulated proteins (FDR cutoff= 0.1).

### Subcellular fractionation

Mouse BMDC were incubated overnight with culture medium or 100 ng/ml upLPS. Human monocytes were partially pre-incubated with 10 µM MCC-950 and stimulated overnight with 100 ng/ml upLPS. Subcellular fractionation was performed using the Cell Signaling Kit (Cell signaling, USA) according to the manufacturer’s instructions. NaK-ATPase was used as a marker for the membrane fraction and GAPDH for the cytoplasmic fraction in immunoblotting.

### Immunoblotting

Cytoplasm and membrane fractions were mixed with 4x Laemmli and β-mercaptoethanol. Equal volumes of fractions were directly loaded in a 4-12% pre-cast gradient gel (BioRad, USA) and separated by SDS-Page. Proteins were transferred to a nitrocellulose membrane (BioRad, USA), blocked with 5% non-fat dietary milk (NFDM, Carl Roth, Germany), and washed and incubated with the primary antibodies were either diluted in 5% NFDM or 5% bovine serum albumin (BSA, Serva, USA) overnight at 4°C: hIL-1α (1:500, Santa Cruz, USA), mIL-1α (1:1,000, R&D, USA), GAPDH (1:5,000, Santa Cruz, USA), NaK ATPase (1:10,000, Abcam, United Kingdom). The membrane was incubated with the secondary antibody coupled with horse radish peroxidase diluted in 5% NFDM for 1h on the following day. Classico Western HRP Substrate (Millipore) or SuperSigna West Femto (Thermo Fisher Scientific, USA) were used for development on iBright 1500 (Thermo Fisher Scientific, USA).

### Proximity Ligation Assay (PLA)

Duolink In Situ Red Starter Kit Mouse/Rabbit (Sigma Aldrich, USA) was used and performed following the manufacturer’s instructions. Briefly, HUVEC cells were seeded on a µ-slide Angiogenesis (ibidi, Germany) and were partially pre-stimulated with neutralizing IL1R1 antibody (1:100, R&D Systems, USA) for 1h. Unstimulated or upLPS-stimulated monocytes were then added for 4h per well. Monocytes were washed 4 times to eliminate residual upLPS before seeding on HUVEC cells. After 4h, the supernatant was aspirated carefully, and cells were fixed with ROTI Histofix 4 % (Carl Roth, Germany) for 20 min at room temperature. The primary antibodies hIL-1α (1:50, Santa Cruz, USA) and IL1R1 Polyclonal Antibody (1:100, Invitrogen, USA) were diluted in Duolink antibody dilution and stained overnight. 1 µg/ml rabbit IgG (Dianova, Germany) was included as an isotype control.

### Monocyte adhesion assay

To determine monocyte-to-endothelial adhesion, HUVEC cells were seeded on 96-well Flat Clear Bottom Black Polystyrene Microplates (Corning, USA). Isolated primary monocytes were stimulated with upLPS overnight or kept in culture medium. The next day, stimulated monocytes were partially pre-incubated with a neutralizing IL-1α antibody (1:1,000, Invivogen, USA) for 1h. Afterward, monocytes were labeled with CellTrace™ Calcein Red-Orange, AM (Invitrogen, USA) for 30 min at 37° C. Monocytes were washed 4 times to eliminate residual upLPS and were co-incubated with HUVECs. After 4 h, fluorescence intensity was measured, as well as the remaining intensity after up to 4 washing steps. %adhesion of monocytes was calculated as stated: ((F_remaining fluorescence_ - F_blanc_)/(F_total_- F_blanc_))x100.

### Flow cytometry analysis d of csIL-1α and VCAM1

For the staining of cell-surface IL-1α ^12^, cells were harvested and washed once with PBS. The cell number was adjusted to 1×10^6^/ml, and cells were resuspended in FACS buffer (1% BSA, 0.05 % sodium azide in PBS). Trustain mouse Fc block (1µg/ml, Biolegend, USA) was added and incubated for 10 min on ice. Anti-biotin IL-1α antibody (1µg/ml, Biolegend, USA) or rat IgG isotype control (1 µg/ml, Biolegend, USA) was added and incubated for 30 min on ice. Anti-Streptavidin PE was added (2,5 µg/ml, eBioscience, USA) and incubated for 30 min on ice in the dark. Cells were incubated with 5 µl of 7AAD (BD Bioscience, USA) for 10 min at room temperature; cells were immediately measured. For the assessment of total IL-1α, cells were fixed after harvest with 250 µl of Cytofix (Biolegend, USA) for 20 min at 4° C. Staining of IL-1α was performed as described, but FACS buffer was exchanged with Permeabilisation buffer (Biolegend, USA). To detect VCAM1 on HUVECs, cells were partially pre-stimulated with neutralizing IL1R1 antibody (1:100, R&D Systems, USA) for 1h. Unstimulated and upLPS-stimulated monocytes were added for 4h. Tnfα served as a positive control. Monocytes were washed 4 times to eliminate residual upLPS before seeding on HUVEC cells. Up-LPS stimulated HEK293 and handled as stimulated monocytes and were included as a control to verify the successful elimination of LPS (Figure S3A). After 4h, the supernatant was discarded, and cells were fixed with 0.5% Roti-Histofix (Carl Roth, Germany) and harvested. Staining was performed using 5 µg/ml anti-human CD106 APC Antibody (Biolegend, USA) or 5 µg/ml mouse IgG2a kappa Isotype control (eBM2a) APC (Thermo Fisher Scientific, USA) as the isotype control, for 45 min on a rotator at 4°C. Cells were measured at the BD FACS Lyrica. FlowJo® Software (Version 10.8.1) was used for further analysis.

### Myristoylation assay

Myristoylation was detected using a myristoylated protein assay kit (abcam, USA) following the manufacturer’s instructions. Cells without myristic acid labeling served as background controls. To test the myristoylation inhibitor, cells were pre-incubated with 1 µM n-myristoyltransferase inhibitor IMP-1088 (Cayman Chemical, USA) for 1h before stimulation with 100 ng/ml upLPS overnight.

### ELISA

Human (R&D Systems, USA) and mouse IL-1α (R&D Systems, USA) DuoSet ELISA kits, as well as human (R&D Systems, USA) and mouse IL-1β (R&D Systems, USA) DuoSet ELISA kits, were used in combination with DuoSet ELISA Ancillary Reagent kit (R&D Systems, USA). The assays were performed following the manufacturer’s instructions.

### Illustrations

Graphical abstract and schematic overviews were generated by using icons from Servier Medical Art, provided by Servier, licensed under a Creative Commons Attribution 3.0 unported license.

### Statistical Analysis

Statistical analyses were performed with GraphPad Prism (version 8; GraphPad Software Inc., La Jolla, CA, USA). Data were tested for a gaussian distribution using the Kolmogorov–Smirnov or D’Agostino-Pearson normality test. Two-tailed unpaired t-test was performed to compare groups if not otherwise stated. One-way ANOVA with Sidak’s multiple comparisons test was performed to compare more than two groups. The significance level was set to p < 0.05.

### Data availability statement

The data underlying this article will be shared on reasonable request to the corresponding author.

## Results

### Il1a^-/-^ mice are protected from hyperlipidemia-induced atherosclerosis, whereas Nlrp3^-/-^ and Il1b^-/-^ mice are not

To investigate the role of IL-1α and the NLRP3 inflammasome in a nongenetic mouse model of atherosclerosis, we induced hypercholesterolemia in WT, Il1a**^-/-^**, Nlrp3**^-/-^**, and Il1b**^-/-^** mice by a single injection of a hyperactive pro-protein convertase subtilisin/kexin type 9 (PCSK9)-adeno-associated virus (rAAV) followed by a Western-type diet (PCSK9-AAV) for 12 weeks (Figure 1A). PCSK9-AAV control mice develop hypercholesterolemia and spontaneous atherosclerosis (Figure 1B-H), as recently described by our group ^15^,. Cholesterol levels were similar in PCSK9-AAV and Il1a, Nlrp3, and Il1b deficient mice (Figure 1B). Il1a*^−/−^* mice showed reduced atherosclerotic plaque area by 61.3 ± 4.6 % (Figure 1C, D) associated with lower lipid accumulation (Oil-red O, Figure 1E, F) compared to PCSK9-AAV control mice (1.9 ± 1.22 % in Il1a*^−/−^* vs. 5.2 ± 2.4 % in PCSK9-AAV mice). The development of atherosclerosis in Nlrp3^−/−^ and Il1b^−/−^ mice did not differ from the control wildtype animals (Figure 1C-F). No significant difference in macrophage infiltration in plaques (MOMA-2 positive cells) was observed in Il1a*^−/−^*, Il1b*^−/−^*, and Nlrp3*^−/−^* mice compared to PCSK9-AVV animals (Figure 1G, H).

### Il1a^-/-^, Nlrp3^-/-^ and Il1b^-/-^ mice show reduced levels of circulating cytokines

To determine whether the atheroprotective effect in Il1a^-/-^ mice are associated with changes in the serum protein profile, we used the Olink Target 96 Mouse Exploratory Panel. The 92 proteins measured encompass various biological processes and thus provide an overview of regulated signaling pathways. The Serum of PCSK9-AAV control mice showed increased proteins annotated to cell-activating signaling pathways and vasculature-regulating mechanisms (Figure 2A, B). Il1a^-/-^ led to a significant downregulation of pro-inflammatory proteins, such as IL-6, IL-1β, and CCL-2, compared to PCSK9-AAV (Figure 2C). IL-6 was also downregulated in Nlrp3^-/-^ and Il1b^-/-^ but both groups show similar plaque development to PCSK9-AAV (Figure 1C). Normal chow control group (NC) and Il1a^-/-^ animals did not share similar regulated proteins compared to PCSK9-AAV (Figure 2D). Even though IL-1β was significantly upregulated in PCSK9-AAV group, serum IL-1α levels did not show a significant difference between NC and PCSK9-AAV (Figure 2E).

**Figure 2:**
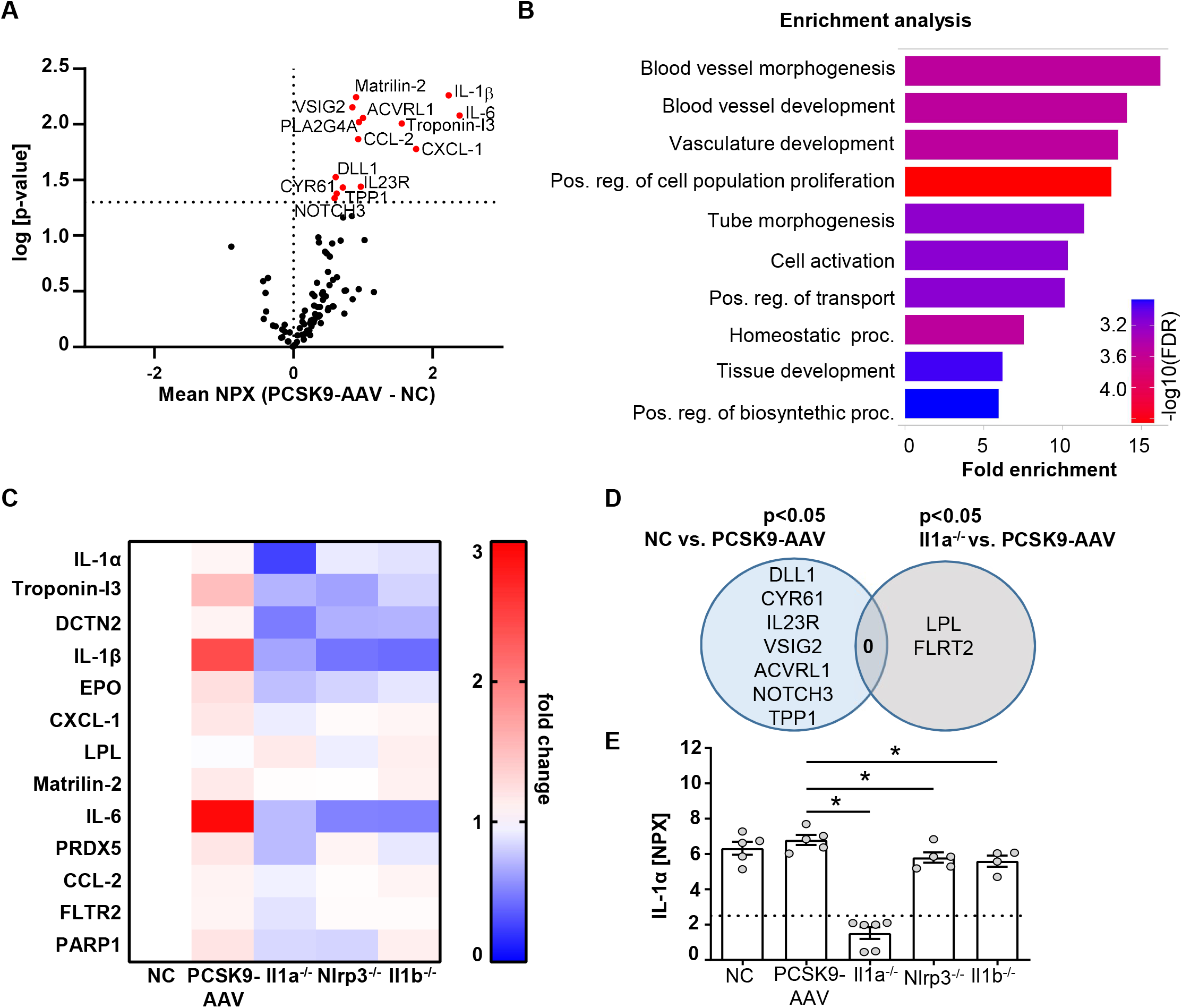
Il1a^-/-^, Nlrp3^-/-^, and Il1b^-/-^ mice show reduced levels of circulating cytokines. **A:** Volcano plot of proteins regulated in Olink mouse exploratory panel comparing serum protein levels of mice fed a NC vs. PCSK9-AAV mice. Red data points indicate the difference of the NPX mean between PCSK9-AAV and NC for each protein. The dashed line intersecting the y-axis indicates the significance of p<0.05. **B:** Shiny GO ^35^ (Version 0.77) enrichment analysis annotating the significantly regulated proteins to the Gene Ontology (GO) Biological Process (false detection rate (FDR) cutoff: 0.1). The top 10 regulated pathways are presented as a barplot with the colors indicating the -log10 FDR, with red as the highest and blue as the lowest. **C**: Heatmap depicting the dynamics of proteins significantly regulated in Il1a^-/-^ compared to PCSK9-AAV. Data is presented as fold change of PCSK9-AAV, Il1a^-/-^, Nlrp3^-/-^, and Il1b^-/-^ to NC. **D:** Venn Diagram representing serum proteins specifically regulated in only NC or Il1a^-/-^ animals compared to PCSK9-AAV. Shared regulated proteins are displayed at the intersection of both areas. **E:** Bargraphs of serum IL-1α levels measured in the Olink 96 mouse exploratory panel. The dashed line indicates the limit of detection of the assay. Data is presented with the Olink NPX value as mean ± SEM, *p<0.05. PCSK9-AAV8 (n=5), Il1a^-/-^ (n=6), Nlrp3^-/-^ (n=5), and Il1b^-/-^ (n=4).

### Cell surface translocation of IL-1α is not influenced by NLRP3 and IL-1β in murine BMDC

Since serum levels of IL-1α were not increased in atherosclerotic PCSK9-AAV mice (Figure 2E), we investigated the membrane-bound form of IL-1α. The shuttle from the cytoplasm to the plasma membrane requires de-novo synthesis of IL-1α, mediated by the TLR/NFκB signaling pathway. BMDCs, as an important representative of circulating immune cells, were used to study the translocation of IL-1α from the cytoplasm to the plasma membrane after a single stimulus of 100 ng/ml ultrapure LPS (upLPS) and cell fractionation (Figure 3A). Stimulation of the TLR leads to the accumulation of pro-IL-1α in the plasma membrane fraction (Figure 3B, C). The purity of the cell fractions was confirmed with markers for cytoplasm (glyceraldehyde-3-phosphate dehydrogenase (GAPDH)) and for plasma membrane (sodium-potassium ATPase (NaK-ATPase)). GAPDH was not detected in the membrane fraction, confirming no contamination of cytoplasmic IL-1α. The translocation to the plasma membrane was validated by flow cytometry on stimulated, unfixed cells (Figure 3D). 7-AAD was used to discriminate between dead and live cells and analyze the cell-surface IL-1α only in unfixed 7-AAD negative cells. (Figure 3D). LPS increased de-novo synthesis of total IL-1α from 4.9 ± 6.6 % to 42.9 ± 7.6 % (fixed cells, Figure 3E) and induced the translocation of IL-1α to the plasma membrane (unfixed 7-AAD-cells, Figure 3F). LPS did not increase the permeability of the cells (Figure 3G). FACS analysis demonstrated that 5.3 ± 1.9 % of the cells showed IL-1α plasma membrane localisation after LPS stimulation (Figure 3F). Stimulated BMDCs of PCSK9-AAV, Nlrp3^-/-^, and Il1b^-/-^ expressed IL-1α in their cytoplasm, which was translocated to the plasma membrane upon LPS stimulation (Figure 3H, I). Even though LPS alone was sufficient for the translocation of IL-1α to the plasma membrane, BMDCs did not secrete IL-1α or IL-1β to the extent of the positive control (LPS+ ATP, Figure S2B, C), which is in line with our *in vivo* findings.

**Figure 3:**
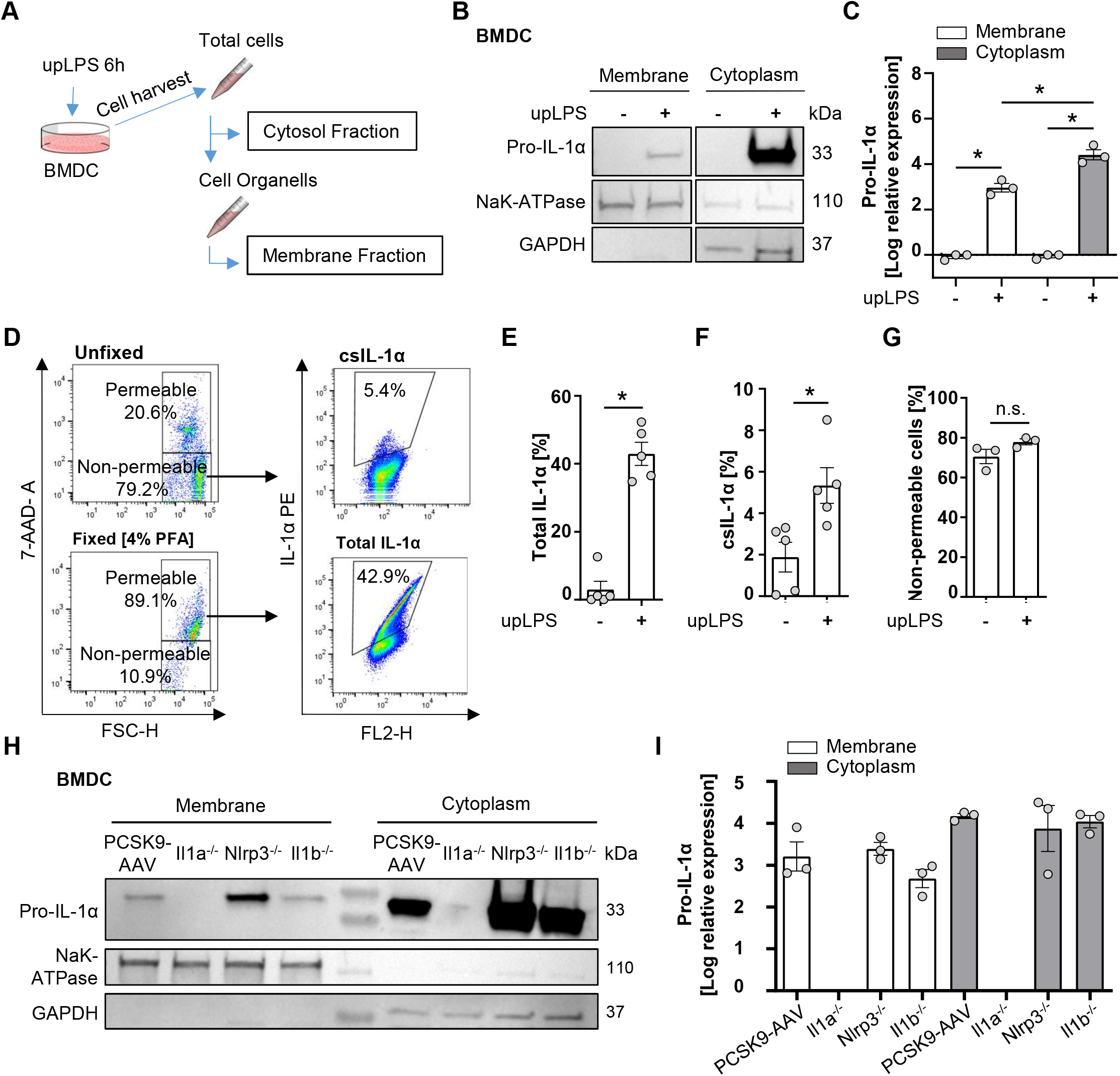
Cell surface translocation of IL-1α is not influenced by NLRP3 and IL-1β in murine BMDC. **A:** Schematic overview of the experimental setup. Bone marrow derived dendritic cells (BMDCs) were replated after 7 days of differentiation and stimulated with 100 ng/ml ultrapure Lipopolysaccharide (upLPS) for 6 hours. Afterward, cells were harvested, and fractions were isolated using a detergent-based method. **B**: Representative immunoblot of the membrane and cytoplasmic fraction of BMDCs with or without upLPS stimulation. Protein levels of IL-1α, GAPDH (glycerinaldehyde-3-phosphate-dehydrogenase, cytoplasmatic marker), and NaK ATPase (sodium– potassium ATPase, membrane marker) are presented. **C:** Densiometric quantification of IL-1α in cytoplasmic fraction normalized to GAPDH and IL-1α in membrane fraction normalized to NaK ATPase. **D:** Representative flow cytometry scatter plot of staining for IL-1α on stimulated BMDCs after gating for viable, non-fixated (7AAD-) cells. Total IL-1α was measured on fixed cells. **E:** Barplot depicting the percentage of total IL-1α positive monocytes (7AAD+) with or without upLPS stimulation. **F:** Barplot depicting the percentage of csIL-1α positive monocytes (7AAD-) with or without upLPS stimulation. **G:** Barplot representing the percentage of non-permeable monocytes with or without upLPS stimulation. **H:** Representative immunoblot of membrane and cytosolic fraction from upLPS-stimulated BMDCs of PCSK9-AAV8, Il1a^-/-^, Nlrp3^-/-^, and Il1b^-/-^ animals. Protein levels of IL-1α, GAPDH (cytoplasmatic marker), and NaK ATPase (membrane marker) are presented with corresponding densiometric quantification (**I**) of log-transformed IL-1α expression. All data are presented as mean ± SEM, *p<0.05.

### IL-1α surface expression on human monocytes induces IL1R1-mediated VCAM1 expression and monocyte adhesion on endothelial cells

To confirm the interaction of monocytic csIL-1α with endothelial IL1R1, a close proximity ligation assay (PLA) was performed. Incubation of LPS-stimulated monocytes with primary human endothelial cells (HUVEC) resulted in a positive PLA fluorescence signal which was not observed under control conditions (Figure 4A). To validate the pro-atherogenic effect of csIL-1α, VCAM1 expression on HUVECs was measured after treatment with LPS-stimulated monocytes. Stimulated monocytes increased VCAM1 expression compared to treatment with unstimulated monocytes (25.9 ± 7.1 % vs. 2.5 ± 1.4 % VCAM pos. cells) on HUVECs, which was significantly reduced by the administration of neutralizing IL1R1 antibody (18.6 ± 6.3% VCAM1 pos. cells) (Figure 4B, C) indicating that csIL-1α/IL1R1 binding and signaling plays a major role in VCAM1 expression and can be abrogated by blocking of IL1R1. In addition, the induced VCAM1 expression through csIL-1α/IL1R1 binding promoted the adhesion of monocytes to HUVECs. Therefore, calcein-labeled monocytes were incubated with HUVECs, and the remaining fluorescence after washing was measured (Figure 4D). Monocytes pre-stimulated with LPS showed more endothelial adhesion than non-stimulated monocytes (6.7 ± 3.6 % vs. 3.5 ± 2.5 %). To study whether monocyte adhesion is dependent on csIL-1α expression and binding on endothelial IL1R1, the cells were co-stimulated with neutralizing IL-1α antibody, which reduced the monocyte adhesion to baseline level (Figure 4E).

**Figure 4:**
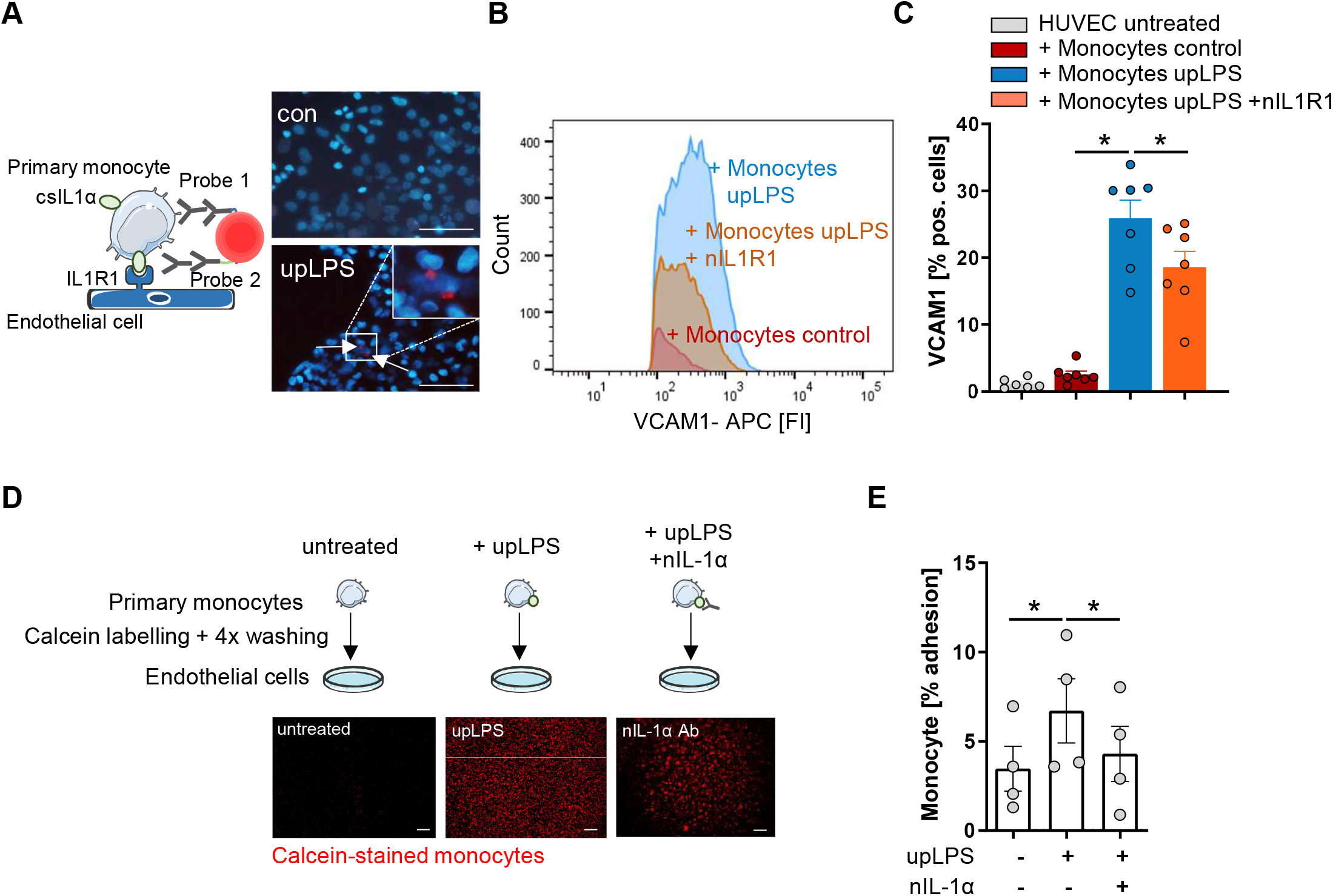
IL-1α surface expression on human monocytes induces VCAM1 expression and leads to increased adhesion on endothelial cells. **A:** Schematic principle of proximity ligation assay (PLA) and representative picture of PLA. Direct binding of csIL-1α to the Interleukin-1 receptor 1 (IL1R1) leads to a fluorescence signal detectable at 594 nm. Human umbilical vein endothelial cells (HUVECs) were treated with monocytes for 6h, stimulated with and without upLPS (100 ng/ml). Cells were imaged at a 40× magnification (scale bar 50µm). HUVECs and monocytes are presented in blue, IL-1α/IL1R1 PLA signal is visible as red dots. **B:** Representative flow cytometry histogram of vascular cell adhesion molecule–1 (VCAM1) stained HUVECs after 4h treatment with upLPS-stimulated monocytes. 10 µg/ml neutralizing IL1R1 (nIL1R1) antibody was added 1h before HUVEC-monocyte co-incubation. **C**: Bar graph depicting the percentage of VCAM1-positive HUVECs after treatment with unstimulated and upLPS-stimulated monocytes. HUVECs were incubated with and without nIL1R1 antibody (10 µg/ml) for 1h before co-incubation. Data are presented as mean± SEM of seven independent experiments; *p< 0.05. **D:** Schematic experimental setup of monocyte adhesion assay. Primary monocytes were treated as indicated, labeled with Calcein and 4x washings. HUVECs were treated with and without 100ng/ml neutralizing IL-1α antibody (nIL-1α) 1h before co-incubation. Then, HUVECs were treated with labeled monocytes for 4h. The initial fluorescence of adhering monocytes was measured as well as after two washes. Cells were imaged (4× magnification), scalebar. **E:** Quantification of adhering monocytes to HUVECs presented as mean ± SEM of four independent experiments. Repeated measure ANOVA was performed, followed by Sidak’s multiple comparison test (*p< 0.05).

### Myristoylation regulates csIL-1α translocation in murine bone marrow cells and in human monocytes

The N-terminus of the 31-kDa IL-1α precursor is myristoylated on lysine residues Lys82 and Lys83 ^19^. Bone-marrow cells from the hyperlipidemic atherosclerotic mice (PCSK9-AAV) showed significantly more protein myristoylation than cells from control mice on normal chow (10.0 ± 4.0 % in control vs. 76.0 ± 35.7 % in PCSK9-AAV mice) (Figure 5A). Blocking of N-myristoyltransferases 1 and 2 (NMT1/2) with IMP-1088 [1µM] reduced protein myristoylation by 62 ± 12.9 % (p< 0.05) (Figure 5B) in human monocytes under control conditions. Pre-incubation of human monocyte cells with IMP-1088 [1µM] for 1h before LPS treatment reduced the cell surface expression of IL-1α by 35 ± 8.7 %, p<0.05 (Figure 5C).

**Figure 5:**
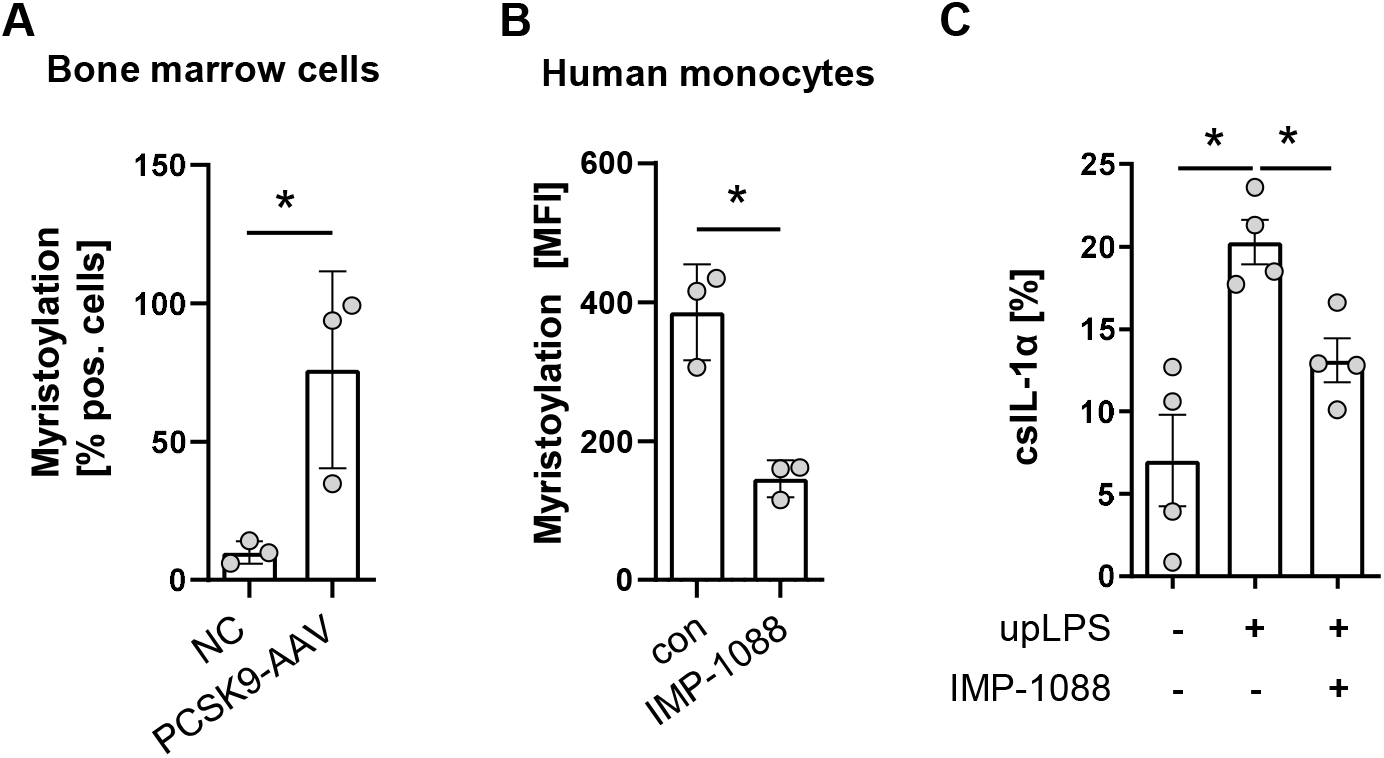
Myristoylation regulates csIL-1α in murine bone marrow cells and human monocytes. **A:** Barplot depicting the percentage of murine bone marrow cells with myristoylated proteins. Bone marrow cells were cultured and myristoylated proteins were labeled overnight. Data are presented as mean ± SEM of NC (n=3) and PCSK9-AAV (n=3), *p<0.05. **B:** Mean fluorescence intensity of human primary monocytes under culture conditions (con) or with overnight incubation of N-myristoyltransferase inhibitor IMP-1088 [1 µM]. Data are presented as mean ± SEM of three independent experiments. One-sided paired t-test was performed, *p<0.05. **C:** Percentage of csIL-1α presenting monocytes stimulated with 100 ng/ml upLPS and 1 µM IMP-1088 as indicated. Data are presented as mean ± SEM of four independent experiments, *p<0.05.

## Discussion

The present study provides evidence for a significant contribution of IL-1α during the development of atherosclerosis. Our novel, nongenetic model of atherosclerosis revealed that circulating pro-inflammatory cytokines are not the only driving factor in atherogenesis. At the observed time points in early atherogenesis, IL-1α at the cell surface mediates leukocyte adhesion to endothelial cells independently of the Nlrp3 inflammasome.

Cytokines are an important factor in immune cell activation and mediators of sterile inflammation. The study demonstrates that pro-inflammatory proteins such as IL-1β, IL-6, and CCL-2 are upregulated in the serum of PCSK9-AAV animals (Figure 2D-F). However, Nlrp3^-/-^ and Il1b^-/-^ animals showed significantly reduced protein levels of IL-1β and IL-6 compared to PCSK9-AAV despite similar plaque areas. The novel observation is that plaque development after 12 weeks of a high-fat diet appears to be largely independent of circulating pro-inflammatory cytokines.

Interestingly, circulating IL-1α was not significantly increased in the PCSK9-AAV animals (Figure 2C). However, the deficiency of Il1a led to a reduction of atherosclerotic lesion size, which implies a significant role of the membrane-bound protein in atherogenesis. The data suggest that the protection of Il1a^-/-^ mice from atherosclerosis is at least partially explained by the deletion of IL-1α expression on the surface of monocytes resulting in reduced leucocyte adhesion. Il1a knockout also led to significant downregulation of CCL-2 and CXCL-1 (Figure 2C, D). This finding highlights the role of IL-1α in monocyte adhesion, not only by upregulation of VCAM1 (Figure 2B, C) but also by reduction of chemokines involved in the transmigration of monocytes into the subendothelial space ^20^. Il1a^-/-^ animals also show a reduction in circulating IL-1β (Figure 2C, dashed line). These results are consistent with previous publications ^21,22^. However, although Il1a^-/-^ animals show reduced IL-1β secretion, this appears not to be the main protective factor from atherosclerotic development since Il1b^-/-^ animals show no reduction in plaque size.

Chen et al. provided a mechanism for IL-1α surface presentation via IL1R2 and GPI anchors ^12^. They showed that Il1r2 deficiency in mice reduced levels of csIL-1α, but the mechanism remained unclear. Our data show that csIL-1α is reduced in primary monocytes by adding the myristoylation inhibitor IMP-1088 before stimulation (Figure 5C). Protein N-myristoylation is an essential fatty acylation catalyzed by N-myristoyltransferases (NMTs), which is vital for proteins participating in various biological functions, including signal transduction, cellular localization, and oncogenesis ^23^. Lately, myristoylated IL-1α was linked to the mitochondrial membrane by binding to cardiolipin. The myristoylation at the lysine residues 82 and 83 increased the association between proteins and lipids and, thus, the integration of IL-1α in the membrane lipid bilayer in the presence of Ca^2+^ ^24^. CsIL-1α levels do not return to baseline, indicating that the different mechanisms of tethering IL-1α to the membrane are essential for the presentation, even though the exact mechanism of transport from IL-1α to the membrane remains elusive. Metabolic stimuli and TLR ligands can induce csIL-1α expression on myeloid cells, making this cytokine a considerable candidate in atherosclerosis initiation, driven by high levels of circulating free fatty acids or other metabolic stimuli, such as cholesterol or high glucose ^25^.

Nlrp3 deficiency did not reduce atherosclerotic lesion size compared to PCSK9-AAV animals in the presented data. Previous Nlrp3^-/-^ studies are controversial and differ in their genetic background and diet composition. Using ApoE^-/-^ mice crossed with either Nlrp3^-/-^, Casp1^-/-^ or Asc^-/-^, mice did not show a reduction in plaque size, macrophage infiltration, or plaque stability after 11 weeks of high-fat diet compared to ApoE^-/-^ mice ^4^. However, chimeric Ldlr^-/-^ mice transplanted with bone marrow from Nlrp3^-/-^, Asc^-/-^, and Il1a^-/-^ mice revealed reduced plaque area after 4 weeks of a high-fat diet ^26^. The PCSK9-AAV8 atherosclerosis model used in this study has the advantage of being independent of genetic alterations to induce atherosclerosis. Knockout of Nlrp3 results in a reduction in circulating cytokines but not in plaque size, lipid accumulation, or macrophage infiltration. Thus, the study contributes to the observation that a constitutive Nlrp3 knockout does not protect against the development of atherosclerotic plaques.

Previous studies have shown that csIL-1α has an implication in different pathologies and is tested as a therapeutic target in different diseases. Monocytes from patients with acute myocardial infarction (AMI) and chronic kidney disease (CKD) exhibit increased levels of csIL-1α. The expression of csIL-1α was associated with an increased risk for atherosclerotic cardiovascular disease events ^20^. Bermekimab (MABp1) is a human antibody targeting IL-1α and was used in patients with refractory cancer. It was well tolerated and led to a decrease in plasma IL-6 ^27^. The antibody was also assessed in phase II clinical study with patients suffering from type II diabetes mellitus. MABp1 improved glycemia and reduced C-reactive protein, an important prognostic marker of systemic inflammation ^28^. Anakinra, an IL1R antagonist, has already been approved by the FDA and is beneficial for patients with rheumatoid arthritis and autoimmune diseases ^29^. Treatment of ApoE^-/-^ mice with anakinra reduced plaque size of the aortic arch and serum triglycerides ^30^. However, anakinra blocks both IL-1α/IL1R and IL-1β/IL1R signaling, which may exert differential or opposite functions in certain diseases. Due to the diverse nature of IL-1α and its associated functions, it is critical to know precisely which IL-1α isoform to target in immunotherapy. Therefore, it appears highly relevant to uncover the detailed role of IL-1α isoforms and their regulation (e.g., by myristoylation) in various diseases.

Knockout of Il1b did not affect lesion size in the PCSK9-AAV8 model of hypercholesterolemia (Figure 1D). However, other studies show a plaque size reduction in ApoE^-/-^/Il1b^-/-^ animals ^31,32^. These controversies could be related to the animal model. Ldlr-deficient animals exhibit significantly smaller lesion sizes and necrotic cores than ApoE^-/-^ mice at similar time points ^33^. In addition, Kamari et al. pointed out that Il1b^-/-^ leads to only 32% plaque reduction, whereas Il1a^-/-^ reduces plaque size by 52% in ApoE double knockout animals ^32^. Vromman et al. investigated the effects of monoclonal antibodies against IL-1α and IL-1β in atherosclerosis *in vivo* which demonstrated that IL-1β has a profound effect on late-stage atherosclerosis by increasing IL-10 in the plasma. In contrast, IL-1α appeard to be more critical in early atherosclerosis due to its influence on arterial outward remodelling ^34^. Therefore, the effect of Il1b-knockout might not be as pronounced in the PCSK9-AAV8 model compared to ApoE^-/-^ animals but might be detectable at a later time point exhibiting progressed atherosclerotic lesions.

As a limitation, our study characterized only one-time point. Our previous study observed that the inflammatory cytokines such as TNF-α increase significantly when comparing the AAV8-PCSK9 model at 12 weeks vs. 20 weeks ^15^. IL-1β may have a greater impact on atherosclerosis in established plaques. Therefore, it would be interesting to examine the changes in secretome as well as plaque characteristics in the applied knockout animals in combination with the PCSK9-AAV8 model at later time points in the future.

In conclusion, IL-1α deficiency reduces atherosclerotic plaque development. Low levels of pro-inflammatory cytokines in the knockout animals indicated independence of plaque development from circulating cytokines at this time point of the disease since csIL-1α mediates initial leucocyte-to-endothelial adhesion and activation of endothelial cells. Importantly, these data highlight the importance of IL-1α in atherosclerosis and the need for detailed understanding of the mechanisms of the translocation and presentation of IL-1α to the plasma membrane as a target for novel therapies.

## Acknowledgment

We thank Anja Barnikol-Oettler, Andreas Hauck, Ellen Becker, and Ihsan Gadi for their excellent technical support.

