## Supplementary Figures for "Membrane-bound Interleukin-1α mediates leukocyte adhesion during atherogenesis"

### Supplements
#### Figure S1

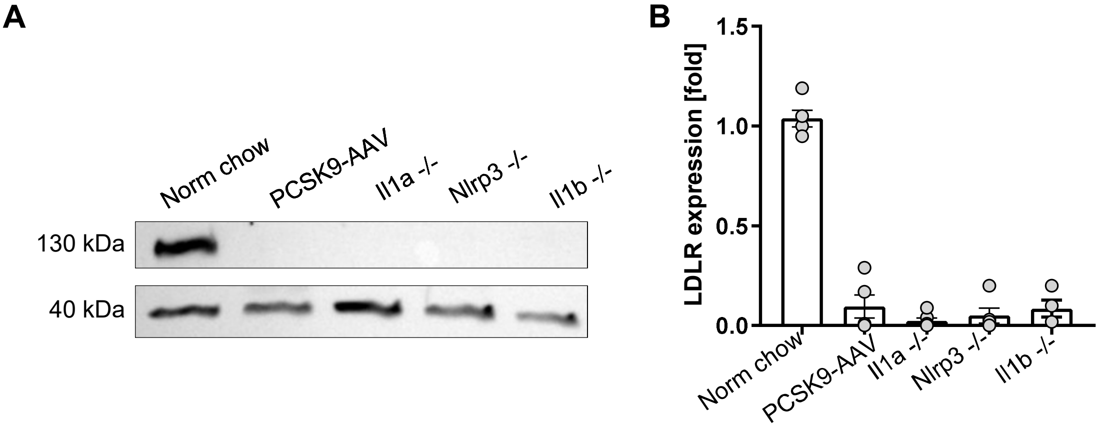


**Supplementary Figure S1:**

**A**: Representative immunoblot of the LDL-receptor (LDLR) expression and β-actin expressed in liver tissue
**B:** Densiometric quantification of LDLR normalized to β-actin.

**Supplements**
**Figure S2**

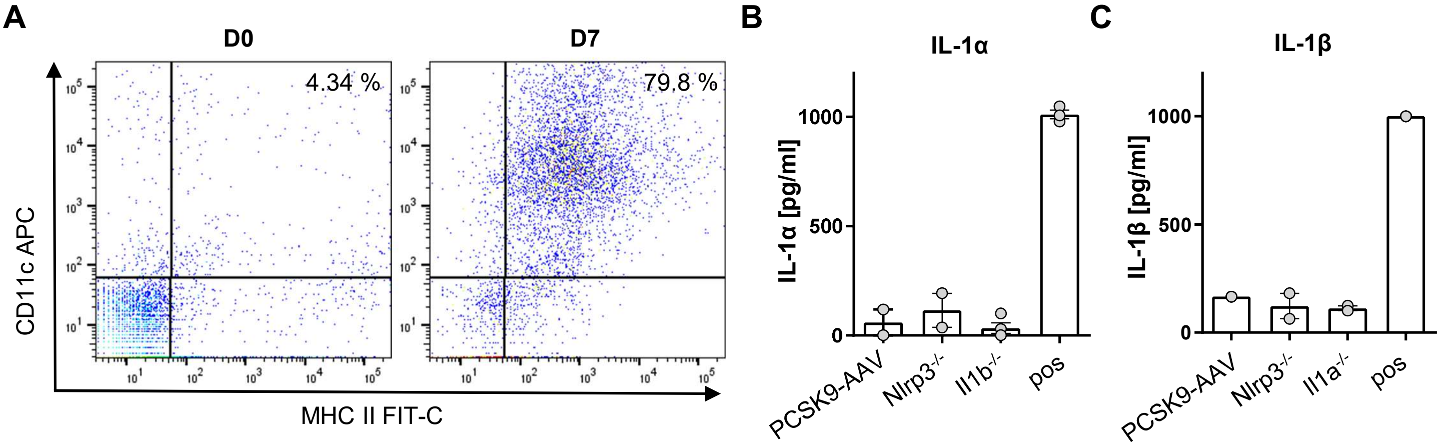


**Supplementary Figure S2:**

**A**: Representative flow cytometry scatter plot of staining for double positive BMDCs for CD11c and MHC II on BMDCs at the first day of differentiation (D0) and 7 days of differentiation (D7). **B:** Secreted IL-1α after 6h of upLPS stimulation in the supernatant of PCSK9-AAV, Nlrp3$^{-/-}$ and Il1b$^{-/-}$. UpLPS (100ng/ml) stimulation with 5mM ATP stimulation for the last 30 min served as the positive control (pos). **C:** Secreted IL-1α after 6h of upLPS stimulation in the supernatant of PCSK9-AAV, Nlrp3$^{-/-}$ and Il1b$^{-/-}$. UpLPS (100ng/ml) stimulation with 5mM ATP stimulation for the last 30 min served as the positive control (pos).

**Supplements**
**Figure S3**

**A**

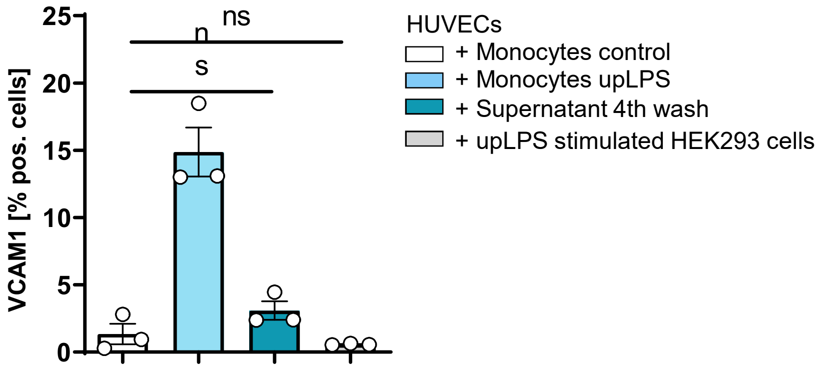


**Supplementary Figure S3:**

**A**: Bar graph depicting the percentage of VCAM1- positive HUVECs after treatment with unstimulated and upLPS-stimulated monocytes. Supernatant after the 4$^{th}$ wash, as well as upLPS, stimulated HEK293 cells and served as the negative control. Data are presented as mean± SEM of three independent experiments; *p< 0.05.
